# TRESK background potassium channel in MrgprA3^+^ pruriceptors regulates acute and chronic itch

**DOI:** 10.1101/2024.01.25.577205

**Authors:** Júlia Llimós-Aubach, Alba Andres-Bilbe, Anna Pujol-Coma, Irene Pallás, Josep Maria de Anta, Concepció Soler, Núria Comes, Gerard Callejo, Xavier Gasull

## Abstract

TRESK (K2P18.1) is a background K^+^ channel expressed in sensory neurons, where it modulates the resting membrane potential, action potential firing and neuronal excitability. A subset of these sensory neurons, which express specific TRPs and Mas-related G protein-coupled receptors (Mrgprs), are activated by pruritogens and mediate itch sensations. Because TRESK is involved in somatosensation and pain transduction, we evaluated the contribution of this channel to pruritic sensitivity and its potential as a target for the treatment of chronic itch pathologies. By combining RNA in situ hybridization, calcium imaging, electrophysiological and behavioral approaches, we found that TRESK is involved in the modulation of non-histaminergic itch. TRESK is coexpressed with MrgprD^+^ and MrgprA3^+^ in sensory neurons and MrgprA3^+^ neurons from TRESK^-/-^ animals display an enhanced firing compared to WT counterparts. Interestingly, acute itch to intradermal injection of chloroquine is significantly enhanced in the absence of TRESK but not the response to histamine, BAM8-22 or LTC4. TRESK deletion also enhanced chronic itch in mice models of Allergic Contact Dermatitis and Dry Skin. In the mouse model imiquimod-induced psoriasiform dermatitis, the absence of TRESK produced a significantly enhanced scratching behavior, which developed earlier and was more robust. Finally, enhancing TRESK function with the channel activator cloxyquin diminished both acute and chronic itch in WT mice but not in KO animals. In summary, our data indicates that TRESK is involved in regulating the excitability of a subset of sensory neurons that mediate histaminergic-independent itch. Enhancing the channel function with specific activators constitutes a novel anti-pruritic therapeutic method that can be combined with other compounds for the treatment of non-histaminergic itch, for which appropriate treatments are lacking.

## Introduction

Itch is defined as an unpleasant sensation that triggers a desire to scratch (1). Persistent pruritus is associated with several skin diseases (e.g. atopic dermatitis, psoriasis or dry skin) as well as certain systemic diseases (cholestasis, chronic kidney disease or Hodgkin’s lymphoma). Beyond the discomfort it brings, chronic itch severely impairs the quality of life, disrupting sleep and social activities. Moreover, it presents a considerable therapeutic challenge due to the limited availability of treatment options.

Within sensory neurons, a specific subset dedicated to sensing itch, primarily innervating the skin and known as pruriceptors, has been described through both genetic and functional approaches (2,3). It is thought that this system is aimed to remove possible irritants or harmful stimuli from the body surface, thereby preventing further damage. The intensity of pruriceptors’ activation by different stimuli is directly related to the sensation of itch, as the activation of these neurons is the first step in conveying the sensory message to the brain via the spinal cord or the brain stem.

In rodents, three subpopulations of Non-Peptidergic (NP) sensory neurons involved in itch sensing have been described (4): NP1, presenting distinctive expression of Mas-related G-Protein-Coupled Receptor D (MrgprD) and activated by β-alanine; NP2 expressing MrgprA3 and MrgprC11, activated by chloroquine and BAM8-22, respectively; and NP3 expressing histamine and serotonin receptors (5–9). Other itch receptors such as PAR2, interleukins (IL-4, -13, -31 and -33) or thymic stromal lymphopoietin (TSLP) are also expressed in these neurons (3,9–12). In humans, similar populations of pruriceptors have been identified containing the same itch receptors or human orthologs (e.g. MrgprX1 in humans is a functional homolog to MrgprA3 and MrprC11 in rodents). Nevertheless, the difference between peptidergic and non-peptidergic sensory neurons subtypes seems not to be present in humans, and substantial overlapping between neuronal markers is present, thus pruriceptors also express several neuropeptide subtypes (13,14).

Beyond the membrane receptors activated by pruritogens, different ion channels such as TRPV1, TRPA1 or ANO1 appear to play a role in the transduction pathway, leading to neuronal depolarization and initiation of action potentials firing (10,12,15,16). Interestingly, in MrgprA3-expressing neurons, different transduction pathways can induce either itch or pain, depending on the activation of metabotropic or ionotropic membrane receptors, respectively (17). In addition, gain-of-function mutations on Nav1.9 (SCN11A) cause a pruritic phenotype in mice and severe pruritus in humans, which has been associated with its high expression in itch nociceptors (18). These findings suggest that each neuronal subtype presents a characteristic pattern of ion channel expression, defining their intrinsic physiological properties, activation mechanism, action potential characteristics, and firing pattern (19).

Two-pore domain potassium channels (K_2P_) are expressed in different sensory neuron subpopulations, including nociceptors, where they predominantly carry “leak” or background hyperpolarizing current (20). Their electrophysiological properties allow them to carry K^+^ currents across a broad range of membrane potentials, positioning them as crucial regulators of neuronal excitability. This is achieved by decreasing the likelihood of depolarizing stimuli reaching action potential threshold and shaping the neuron firing response (20). In both humans and rodents, TRESK (<u>T</u>WIK-<u>re</u>lated <u>s</u>pinal cord K^+^ channel) exhibit high expression in neurons of the dorsal root ganglia (DRG) and trigeminal ganglia (TG), where it is particularly enriched compared to other neural and non-neural tissues (4,21–24). We previously characterized the role of TRESK in nociception, providing further insights on the specific functions of the channel. We have shown that the absence of TRESK expression in knock out animals results in a significant enhancement of mechanical and cold sensitivity, whereas sensitivity to heat remains largely unaffected (24,25). Additionally, mutations in the channel have been linked to migraine pain (26–28).

Analyzing different publicly available transcriptomic studies (4,21,29–32), we noted the selective expression of TRESK K^+^ channel in three distinct subpopulations of sensory neurons, two of them involved in itch transduction (NP1 and NP2). Given its selective expression in these neurons and TRESK electrophysiological properties, we explored the hypothesis that the channel could significantly contribute to the regulation of pruriceptors excitability. Consequently, alterations in its expression or function might result in the persistent activation of pruriceptors, leading to chronic itch. Employing a comprehensive approach involving various techniques, here we establish TRESK as a key regulator in non-histaminergic itch modulation. TRESK coexpression with MrgprD^+^ and MrgprA3^+^ in sensory neurons, along with enhanced firing in MrgprA3^+^ neurons following TRESK deletion, highlights its substantive role. Notably, TRESK deficiency amplifies acute itch triggered by chloroquine and exacerbates chronic itch in diverse murine models. Importantly, manipulating TRESK function with the specific activator cloxyquin emerges as a promising novel approach for anti-pruritic therapy, offering potential for combination with existing compounds to address non-histaminergic itch, an area currently lacking effective treatments.

## Methods

### Animals

All behavioral and experimental procedures were carried out in accordance with the recommendations of the International Association for the Study of Pain (IASP) and were reviewed and approved by the Animal Care Committee of the University of Barcelona and by the Department of the Environment of the *Generalitat de Catalunya*, Catalonia, Spain (#9876, 10191, 10326). Female and male C57BL/6 mice between 8 and 16 weeks old were used in all experimental procedures (RNA extraction and qPCR, in situ hybridization, cell cultures, electrophysiology, calcium imaging and behavior) unless specified otherwise. Mice were kept in a controlled environment at 22°C with unrestricted access to food and water in a 12-hour alternating light and dark cycle.

TRESK (Kcnk18/K2P18.1) knockout mice (KO) and wild-type (WT) littermates were obtained from the KOMP Repository (Mouse Biology Program, University of California, Davis, CA). The TRESK knockout mouse was generated by replacing the complete Kcnk18 gene by a ZEN-UB1 cassette according to the VelociGene’s KOMP Definitive Null Allele Design. At 3 weeks of age, WT or KO newborn mice were weaned, separated, and identified by ear punching. Genomic DNA was isolated from ear snip samples incubated in a solution containing 25mM NaOH and 0,2 mM EDTA (pH 12) at 95°C during 30 minutes, following neutralization with 40 mM Tris-HCl solution at pH 5. Polymerase chain reaction (PCR) was performed with primers to detect the Kcnk18 gene: forward 5’-GGAGAACCCTGAGTTGAAGAAGTTCC-3’ and reverse GGCTCTAACTTTCCTCACTGCACC or the inserted cassette in the KO mice: forward (REG-Neo-F) GCAGCCTCTGTTCCACATACACTTCA and reverse (gene-specific) AGACTTCTCCCAGGTAACAACTCTGC. The PCR mixture contained 9 μl DNA sample, 10 μl Master Mix (Thermo Scientific; Ref. K0171), 0.5 μl (20 μM) forward and reverse primers (final volume of 20 μl). PCR amplifications were carried out with 35 cycles in a programmable thermal cycler (Eppendorf AG, Hamburg, Germany). The program used was: 95°C for 3 min and cycles of 95°C for 30 s, 58°C for 60 s, 72°C for 40 s, with a final extension at 72°C for 5 min. PCR products were analyzed by electrophoresis in 1.5% agarose gels. Once identified, genotyped animals were used as breeders for colony expansion and their offspring were used in all experimental procedures in which mice were required. The transgenic MrgprA3^GFP-Cre^ mouse (Tg(Mrgpra3-GFP/cre)) line was kindly provided by Mark Hoon (NIH) and Xinzhong Dong (Johns Hopkins University, HHMI)(5) and crossed with a Cre-dependent ROSA26^tdTomato^ reporter mouse line (Gt(ROSA)26Sortm14(GAG-tdTomato)Hze) providing the expression and of tdTomato in MrgprA3-expressing neurons in the resulting mouse line (MrgprA3^tdTomato^). This MrgprA3^tdTomato^ mouse line was further crossed with TRESK^-/-^ mice to obtain the MrgprA3^tdTomato^; TRESK^-/-^ mouse line. Primers for CRE recombinase (Cre-F: CAGAGACGGAAATCCATCGC; Cre-R: GGTGCAAGTTGAATAACCGG), Gt(ROSA) WT (forward: GGCATTAAAGCAGCGTATCC; reverse: CTGTTCCTGTACGGCATGG) and Gt(ROSA) Mut (forward: AAGGGAGCTGCAGTGGAGTA; reverse: CCGAAAATCTGTGGGAAGTC) were used. The program for the Cre PCR used was: 95°C for 3 min and cycles of 95°C for 30 s, 53°C for 60 s, 72°C for 30 s, with a final extension at 72°C for 5 min. The program for the Gt(ROSA) PCR used was: 95°C for 3 min and cycles of 95°C for 30 s, 58°C for 60 s, 72°C for 30 s, with a final extension at 72°C for 5 min. Mice generated were used to obtain primary cultures of tdTomato-labeled MrgprA3^+^ sensory neurons from control or TRESK^-/-^ animals for electrophysiological recordings.

### RNAscope in situ hybridization

DRG or TG ganglia were dissected from WT mice and multi-labeling in situ hybridization (ISH) was performed using fresh frozen tissue with the RNAscope^®^ technology (Advanced Cell Diagnostics (ACD), Newark, CA) according to the manufacturer’s instructions. Probes against *mrgpra3*, *mrgprD*, *tubb3* (Tubulin beta 3) and *kcnk18* (TRESK) mRNAs were used in conjunction with the RNAscope^®^ multiplex fluorescent development kit. Images were obtained on a Zeiss LSM880 confocal laser-scanning microscope (Jena, Germany). ImageJ (NIH, Bethesda, MD) was used to analyze images and to determine soma size.

### RNA extraction and quantitative real-time PCR

Mouse tissue samples were obtained from trigeminal ganglia at indicated times, kept in RNAlater solution (Qiagen, Madrid), frozen in liquid N_2_ and stored at −80°C until use. Total RNA was isolated using the Nucleospin RNA (Macherey-Nagel, Germany) and quantified for purity in a Nanodrop^®^ (ThermoFisher Scientific, Waltham, MA, USA). First-strand cDNA was then transcribed using the SuperScript IV Reverse Transcriptase (Invitrogen, ThermoFisher Scientific) according to the manufacturer’s instructions. Quantitative real-time PCR was performed in an ABI Prism 7300 using the Fast SYBR Green Master mix (Applied Biosystems, Waltham, MA, USA) and primers (detailed below) obtained from Invitrogen (ThermoFisher Scientific). Amplification of Glyceraldehyde 3-phosphate dehydrogenase (GAPDH) transcripts was used as a standard for normalization of all qPCR experiments and gene-fold expression was assessed using the ΔC_T_ method. All reactions were performed in triplicate. After amplification, melting curves were obtained and evaluated to confirm correct transcript amplification. Gene-specific primers used were: *tresk* (forward 5′- CTCTCTTCTCCGCTGTCGAG-3′; reverse 5′-AAGAGAGCGCTCAGGAAGG-3′); *mrgprA3* (forward 5′- CTCAAGTTTACCCTACCCAAAGG -3′; reverse 5′- CCGCAGAAATAACCATCCAGAA -3′); *mrgprD* (forward 5′- TTTTCAGTGACATTCCTCGCC -3′; reverse 5′- GCACATAGACACAGAAGGGAGA -3′); *mrgprC11* (forward 5′- ACTCTCTGCTACGGATCATTGA -3′; reverse 5′- TGATTGCTGCATTGCCTAAGATA -3′); *trpa1* (forward 5′- GTCCAGGGCGTTGTCTATCG -3′; reverse 5′- CGTGATGCAGAGGACAGAGAT -3′); *trpv1* (forward 5′- CCCGGAAGACAGATAGCCTGA -3′; reverse 5′- TTCAATGGCAATGTGTAATGCTG -3′); *gapdh* (forward 5′- TCCACTGGCGTCTTCAC -3′; reverse 5′- GGCAGAGATGATGACCCTTTT -3′).

### Primary culture of sensory neurons

Mice were humanely euthanized through cervical dislocation and decapitation under isoflurane anesthesia. The thoracic, lumbar and cervical dorsal root ganglia (DRG) or trigeminal ganglia (TG) were dissected for neuronal culture as previously described (24,33). Briefly, the collected ganglia were maintained in cold (4–5°C) Ca^2+^ - and Mg^2+^ -free Phosphate Buffered Saline solution (PBS) supplemented with 10 mM glucose, 10 mM Hepes, 100 U.I./mL penicillin and 100 μg/mL streptomycin until dissociation. Subsequently, ganglia were incubated in 2 ml HAM F-12 with collagenase CLS I (1 mg/ml; Biochrome AG, Berlin), Bovine Serum Albumin (BSA, 1 mg/ml) and dispase II (5 mg/ml, Sigma) for 1 h 45 min at 37°C followed. Ganglia were then resuspended in culture media (HAM F-12 supplemented with 10% FBS, 100 μg/ml of penicillin/streptomycin and 100 mg/ml of L-glutamine) and mechanical dissociation was conducted with fire-polished glass Pasteur pipettes of decreasing diameters, previously coated with Sigmacote (Sigma) and autoclaved. The resulting neurons were centrifuged at 1000 rpm for 5 min, re-suspended in culture medium, and transferred to 12 mm-diameter glass coverslips pre-treated with poly-L-lysine/laminin. The primary cultures were then incubated at 37°C in humidified 5% CO_2_ atmosphere for up to 1 day before being utilized for patch-clamp electrophysiological recordings or calcium imaging experiments.

### Calcium imaging

Cultured sensory neurons from wild-type and TRESK^-/-^ mice were loaded with 5 μM fura-2/AM (F1221, Invitrogen, ThermoFisher Scientific, Waltham, MA, USA) for 45-60 min at 37°C in culture medium. Coverslips with fura-2 loaded cells were transferred into an open-flow chamber (0.5 ml) mounted on the stage of an inverted Olympus IX70 microscope equipped with a TILL monochromator as a source of illumination. Pictures were acquired with an attached cooled CCD camera (Orca II-ER, Hamamatsu Photonics, Japan) and stored and analyzed on a PC computer using Aquacosmos software (Hamamatsu Photonics, Shizuoka, Japan). After a stabilization period, pairs of images were obtained every 4 s at excitation wavelengths of 340 (11) or 380 nm (12; 10 nm bandwidth filters) in order to excite the Ca^2+^ bound or Ca^2+^ free forms of the fura-2 dye, respectively. The emission wavelength was 510 nm (12-nm bandwidth filter). Typically, 20-40 cells were present in the microscope field (20x). [Ca^2+^]_i_ values were calculated and analyzed individually for each single cell from the 340- to 380-nm fluorescence ratios at each time point. Only neurons that produced a response >10% of the baseline value and that, at the end of the experiment, produced a Ca^2+^ response to KCl-induced depolarization (50 mM) were included in the analysis. Several experiments with cells from different primary cultures and different animals were used in all the groups assayed. The extracellular (bath) solution used was Hank’s Balanced Salt Solution (HBSS): 140 mM NaCl, 4 mM KCl, 1.8 mM CaCl_2_, 1 mM MgCl_2_, 10 mM glucose, 10 mM HEPES, at pH 7.4 with NaOH. Experiments were performed at room temperature. Drugs were bath-applied for 30 s and then washed out. To evaluate the effects of chloroquine on intracellular calcium, Fluo-4 AM was used instead of Fura-2, as this drug has an important light absorption in the UV range, wavelengths used for Fura-2 stimulation. Fluo-4 was stimulated at 494 nm and light emitted at 506 nm was recorded using the same equipment described.

### Electrophysiological recordings

Electrophysiological recordings in sensory neurons were performed as previously described (24,33,34). Briefly, recordings were performed with a patch-clamp amplifier (Axopatch 200B, Molecular Devices, Union City, CA) and restricted to tdTomato-expressing TG neurons from MrgprA3^TdTomato^ or MrgprA3^TdTomato^; TRESK^-/-^ mice, which had a soma diameter <30 μm and largely correspond to nociceptive/pruriceptive neurons (35). Patch electrodes were fabricated in a Flaming/Brown micropipette puller P-97 (Sutter instruments, Novato, CA). Electrodes had a resistance between 2-4 MΩ when filled with intracellular solution (in mM): 30 KCl, 110 K-Gluconate, 2 MgCl_2_, 10 HEPES, adjusted to pH 7.2. Bath solution (in mM): 145 NaCl, 5 KCl, 2 CaCl_2_, 2 MgCl_2_, 10 HEPES, 5 glucose at pH 7.4. The osmolality of the isotonic solution was 310.6±1.8 mOsm/Kg. To study sensory neuron excitability, after achieving the whole-cell configuration of the patch clamp technique, the amplifier was switched to current-clamp bridge mode. Recorded signals were filtered at 2 kHz, digitized at 10 kHz and acquired with pClamp 10 software. Data was analyzed with Clampfit 10 (Molecular Devices) and Prism 10 (GraphPad Software, Inc., La Jolla, CA). Series resistance was always kept below 15 MΩ and compensated at 70-80%. All recordings were done 18-24h after dissociation at room temperature (22-23°C) rather than body temperature to permit direct comparisons to other studies of underlying biophysical mechanisms. Only neurons with a resting membrane voltage below −45 mV were considered for the study. Rheobase was determined with ascending series of 200 ms depolarizing pulses (10 pA increments from −50 pA) while the neuron was held at its resting membrane potential (RMP). Action potential (AP) properties (amplitude, duration, time to AP peak) were measured at the spike fired at the rheobase current. Membrane input resistance (Ir) was measured at the −50 pA current pulse. Neuronal excitability was measured as the number of spikes during a 1s depolarizing ramp from 0 to 500 pA.

### Histology

To evaluate the skin lesions produced by the Imiquimod-induced psoriatic skin model in WT and TRESK^-/-^ animals, a new batch of animals were treated as described previously and sacrificed at day 4. Cheek skin specimens were dissected immediately after animals were sacrificed and fixed in 4% paraformaldehyde in Ca^2+^/Mg^2+^-free PBS and processed for paraffin embedding. 5 μm-thick sections were stained with hematoxylin and eosin. Images were obtained in a bright-field Nikon Eclipse E-800 microscope at 20x and main histological parameters from at least four sections of each mouse were analyzed with ImageJ software (NIH, Bethesda, MD) in a blinded manner by two researchers. In particular, epidermal thickness was determined as the number of epidermal cell layers, hypergranulosis (thickness of the stratum granulosum) was assessed as a percentage of epidermal length with two or more epidermal granular layers, and microabscesses were quantified as the number per epidermal length.

### Behavioral studies

Female and male WT or TRESK^-/-^ mice aged between 8 and 16 weeks were used in all behavioral studies. To mitigate stress-induced variability in the results, mice were acclimated to the experimental room and the experimental setup prior to testing. Behavioral measurements were conducted in a quiet environment, with careful attention to minimize or prevent any distress of the animals. All experimenters were blind to the genotype, or the drug/vehicle assayed.

### Cheek assay of acute itch

To assess acute itch sensitivity, we employed the cheek assay, which allows to differentiate between itch and pain responses to a substance (36). Mice were acclimated to the experimental room for 2 hours one day before testing. Compounds were intradermally injected into the cheek of the animal (10 μL). A characteristic scratching response with the posterior leg is indicative of itch while a wipe with the anterior leg indicates pain, as the animal attempts to eliminate the painful stimulus. The number of scratching bouts towards the injection site during 15-minute period was counted. In experiments using cloxyquin or olive oil (vehicle), animals received an intraperitoneal injection of the drug (50 mg/Kg) 2h before testing.

### Imiquimod-induced psoriasis model of chronic itch

The imiquimod model of psoriatic itch (37) was performed as described, on the mouse cheek. Briefly, on the day before the treatment and under short anesthesia with 2% isoflurane, the mouse’s cheek skin was shaved (0.5 x 0.5 cm). For 7 consecutive days (days 0-6), mice received a daily topical application of 12.5 mg Aldara^®^ cream (5% imiquimod; 3M Pharmaceuticals) on the shaved cheek skin. Control mice was treated similarly with a control vehicle cream (Vaseline Lanette). The behavior of each mouse was recorded during 1h on day 0 (baseline, before applying any treatment), on days 1-6 (before imiquimod re-application) and on day 7. Animals were observed individually in its home cage where they could move freely without any type of restriction. The number of scratching bouts directed to the area were counted as a measure of spontaneous itch.

### Dry skin model of chronic itch

The dry skin model was performed on the mouse cheek as described (38). Briefly, after shaving (0.5 x 0.5 cm), the mouse cheek was treated twice daily during 5 days, with a cotton swab moistened with water (control group) or with a 1:1 mixture of acetone and ether (dry skin group) during 15 s. The treated cheek skin was then washed with water for 30 s. On day 1 (before applying any treatment) and on days 3 and 5 (after the second treatment of the day), the behavior of each mouse was individually observed while the animal freely moved in its home cage. The number of scratching bouts during a 20-min period was counted as a measure of spontaneous itch.

### Allergic contact dermatitis model of chronic itch

To induce allergic contact dermatitis, each mouse was challenged with the contact sensitizer squaric acid dibutyl ester (SADBE; 339792; Sigma, Madrid) to elicit contact hypersensitivity (39). Mice were sensitized by a topical application of 25 μl of 1% SADBE in acetone to the shaved back skin (0.5 x 0.5 cm) once a day for three consecutive days (days 0, 1 and 2). Five days later (day 5), the SADBE-treated group was challenged with a topical application of 1% SADBE to the shaved cheek skin once a day for two consecutive days (days 7 and 8) whereas acetone alone was used as the vehicle control. Spontaneous scratching behavior was observed for 1h on three consecutive days, before the SADBE challenge (day 7) and during the two days after (days 8 and 9).

### Drugs

All reagents and culture media were obtained from Sigma-Aldrich (Madrid, Spain) unless otherwise indicated, including histamine (H7250), β-alanine (146064), chloroquine (C6628), BAM 8-22 (SML0729), cloxyquin (C47000). N-methyl leukotriene C4 (LTC_4_; CAY-13390) was obtained from Cayman Chemical (Ann Arbor, Michigan, United States). All drugs were prepared fresh before every experimental procedure by diluting them in Phosphate Buffered Saline (PBS) at the desired concentration except LTC_4_ that was diluted in PBS from a stock solution in ethanol. Cloxyquin was diluted and injected intraperitoneally in olive oil (40).

### Data analysis

Data are presented as mean ± SEM. Statistical differences between different sets of data were assessed by performing unpaired Student’s t-tests, one-way or two-way ANOVA plus Bonferroni’s or Holm-Šídák’s multiple comparisons tests, as indicated. The significance level was set at p<0.05 in all statistical analyses. Data analysis was performed using GraphPad Prism 10 software and GraphPad QuickCalcs online tools (GraphPad Software, Inc., La Jolla, CA).

## Results

### TRESK colocalizes in sensory neuron subpopulations involved in itch signaling

Single-cell transcriptomic studies of both trigeminal and dorsal root ganglia have revealed predominant TRESK expression in specific sensory neuronal subtypes (4,21,29,30,32,41). The channel is expressed in low-threshold mechanoreceptors involved in touch sensation (NF1; expressing TrkB and Piezo2) but predominantly in non-peptidergic nociceptors subpopulations NP1 and NP2, responsible for itch transduction. To validate these findings, we performed in situ hybridizations and quantified the co-expression percentage of TRESK with known markers of these subpopulations, Mas-related G-protein-coupled receptor D (MrgprD) for NP1 and MrgprA3 for NP2. Trigeminal ganglia slices obtained from 3 different WT mice were assayed for *tubb3*, *kcnk18* (TRESK) and *mrgprA3* or *mrgprD*. From 5852 neurons (Tubb3^+^), 2129 were positive for TRESK^+^ (35.6±6.7%) and 305 for MrgprA3^+^ (4.9±1.5%), mostly being small or medium-sized neurons (Fig 1A). A significant percentage of MrgprA3^+^ neurons (67.9±24.3%, 196 neurons) exhibited a notable degree of TRESK co-expression (Fig 1A,C). Similar results were found in DRGs, where 57.9±4.8% MrgprA3^+^ neurons express TRESK (MrgprA3^+^, 85 neurons, TRESK^+^ 642 neurons, Tubb3^+^, 1865 neurons; Fig 1B,C). TRESK also showed a high percentage of expression in MrgprD^+^ neurons in both TG and DRG neurons, as 60.2±2.4% of MrgprD^+^ trigeminal neurons also express the ion channel (MrgprD^+^, 962 neurons, TRESK^+^ 2789 neurons, Tubb3^+^, 7645 neurons; Fig 1C). In DRGs, the percentage of MrgprD^+^ neurons expressing the channel was slightly higher (73.2±6.2%; MrgprD^+^, 1064 neurons, TRESK^+^ 1509 neurons, Tubb3^+^, 4590 neurons; Fig 1C). Therefore, ISH data confirms transcriptomic studies, showing that TRESK is highly expressed in NP1 and NP2 neuronal subtypes involved in itch transduction.

**Figure 1:**
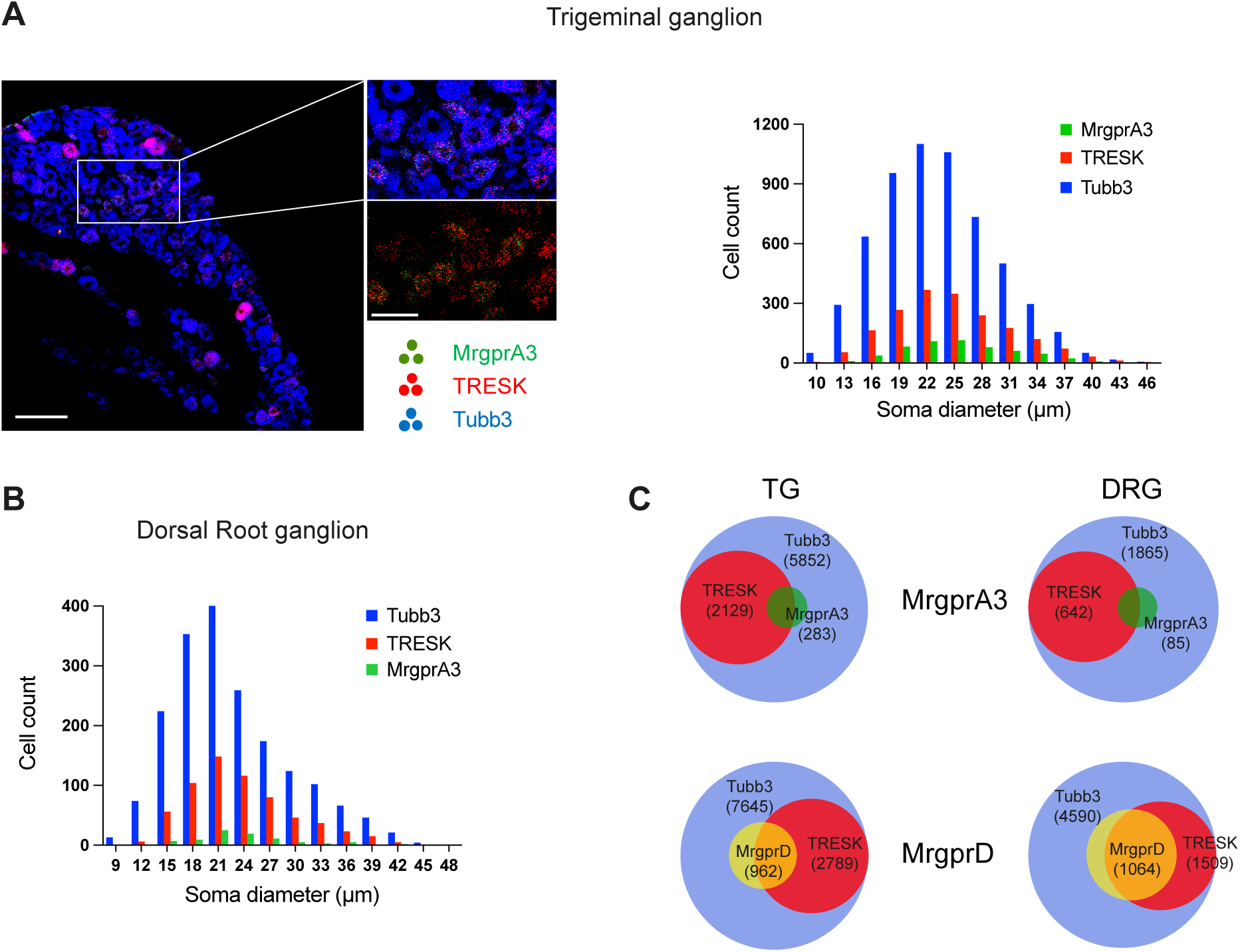
Coexpression of TRESK and Mrg receptors in sensory neurons. **A.** *Left*, Triple-label in situ hybridization of MrgprA3 (green), TRESK (red) and tubulin beta 3 (Tubb3, blue) in the trigeminal ganglion from wild-type mouse. Detailed coexpression of MrgprA3 and TRESK can be seen in neurons (Tubb3^+^) from the inset. *Right*, quantification and distribution of soma size for the three groups, revealing that TRESK and MrgprA3 are predominantly expressed in small and medium sensory neurons. Scale bar: 100 μm; Inset: 50 μm. **B.** Quantification and distribution of soma size for the three groups in dorsal root ganglia sensory neurons showing a similar distribution pattern as observed in trigeminal ganglia. **C.** Venn diagrams depicting the number of cells expressing MrgprA3 and TRESK or MrgprD and TRESK in the total neuronal population (Tubb3^+^) and the overlapping mRNA expression pattern in trigeminal and dorsal root ganglia. Data were obtained from 8-10 sections from n=3-4 mice per ganglia.

### TRESK ablation enhances acute itch

Considering the role of TRESK regulating neuronal excitability (24,26,27,42) and its expression in pruriceptors, we reasoned that the absence of the channel could heighten neuronal activation by pruritogenic compounds. To test this hypothesis, we employed the cheek assay (36) to assess itch responses following the injection of different well-known pruritogens in WT and TRESK^-/-^ mice. As anticipated, chloroquine (CQ) injection (50 μg in 10 μl), a MrgprA3 agonist (43), resulted in significant scratching compared to vehicle injection in both genotypes (Fig 2A). Interestingly, mice lacking TRESK exhibited an enhanced response to chloroquine (WT vs KO: 24.5±13.9 vs. 48.6±26.6 scratching bouts; p<0.01: Fig 2A), suggesting that the absence of the channel facilitates neuronal activation. In contrast, scratching bouts induced by histamine (30 μg; activating H1R receptors), BAM8-22 (50 nmol; activating MrgprC11) and N-met LTC4 (1μg; activating Cysltr2) were similar between WT or KO animals, despite inducing significant scratching in all groups compared to vehicle (Fig 2A). In our hands, cheek injection of β-alanine only produced a mild effect in both WT and KO mice (not shown) despite others have reported significant effects (6). None of these compounds elicited almost any wiping responses (indicative of pain (36)), thus confirming that tested compounds induced itch rather than pain. These results suggest that TRESK modulates specific populations of itch- transducing sensory neurons, but not others, preferentially non-histaminergic itch transduction mediated by MrgprA3^+^ pruriceptors (NP2) activation. To rule out the possibility that the enhanced scratching response in KO mice resulted from an increased number of neurons responding to chloroquine, we performed calcium imaging experiments to determine the percentage of neurons responding to each pruritogen (Fig 2C). No significant differences were found in the number of neurons activated by any of the pruritogens tested or the amplitude of the calcium peaks, indicating that neuronal populations were largely similar and unaffected in KO mice.

**Figure 2:**
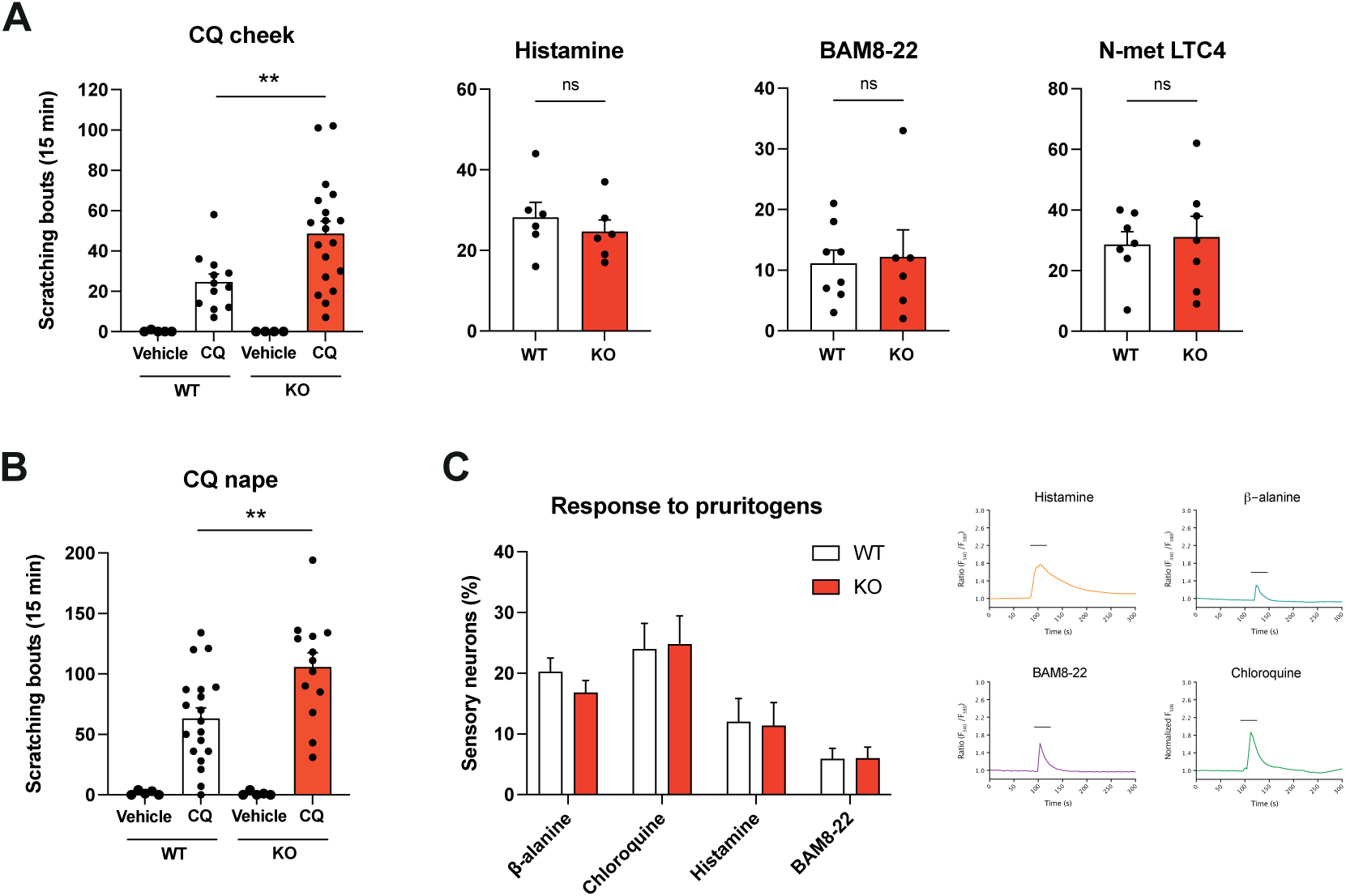
Deletion of TRESK enhances non-histaminergic acute itch. **A.** Intradermal injection of chloroquine (50 μg), histamine (30 μg), BAM8-22 (50 nmol) or N-met LTC4 (1μg) into the cheek induced acute scratching that was significantly greater than vehicle injection. An enhanced response was observed in TRESK^-/-^ animals in response to chloroquine compared to WT mice (**p<0.01, unpaired two-tailed Student’s t-test). No significant differences (ns) were found for the other compounds. **B.** Similar results were obtained when injecting chloroquine in the nape of the neck, a territory innervated by DRG sensory neurons (**p<0.01, unpaired two-tailed Student’s t-test). **C.** *Left*, Percentage of neurons from WT and TRESK^-/-^ mice showing increases in intracellular calcium after challenge with the above-indicated pruritogens (24h after primary culture). Similar percentages of response were obtained in both genotypes. *Right*, Example traces of calcium experiments for each pruritogen tested.

### TRESK regulates MrgprA3^+^ neurons excitability

In a previous study, we observed that TRESK deletion enhanced the excitability of small and medium sensory neurons (24). To specifically test whether TRESK removal enhanced the excitability of MrgprA3+ neurons, we recorded trigeminal neurons from the MrgprA3^tdTomato^; TRESK^+/+^ and MrgprA3^tdTomato^; TRESK^-/-^ mice under current clamp conditions. As shown in figure 3A,B, we did not observe significant differences in the resting membrane potential, input resistance, rheobase, action potential amplitude and width. A significant difference was observed in the time-to-action potential peak, where KO neurons exhibited a shorter time to reach the peak, suggesting a facilitated depolarization phase. Interestingly, when subjected to a depolarizing ramp, KO neurons fired a significant higher number of spikes (5.06±0.96; n=16; p<0.01; Fig 3B,C) compared with their WT counterparts (3.22±0.48; n=46), indicating increased excitability in the absence of TRESK. This suggests that pruritic stimuli might lead to a heightened activation of the pathway.

**Figure 3:**
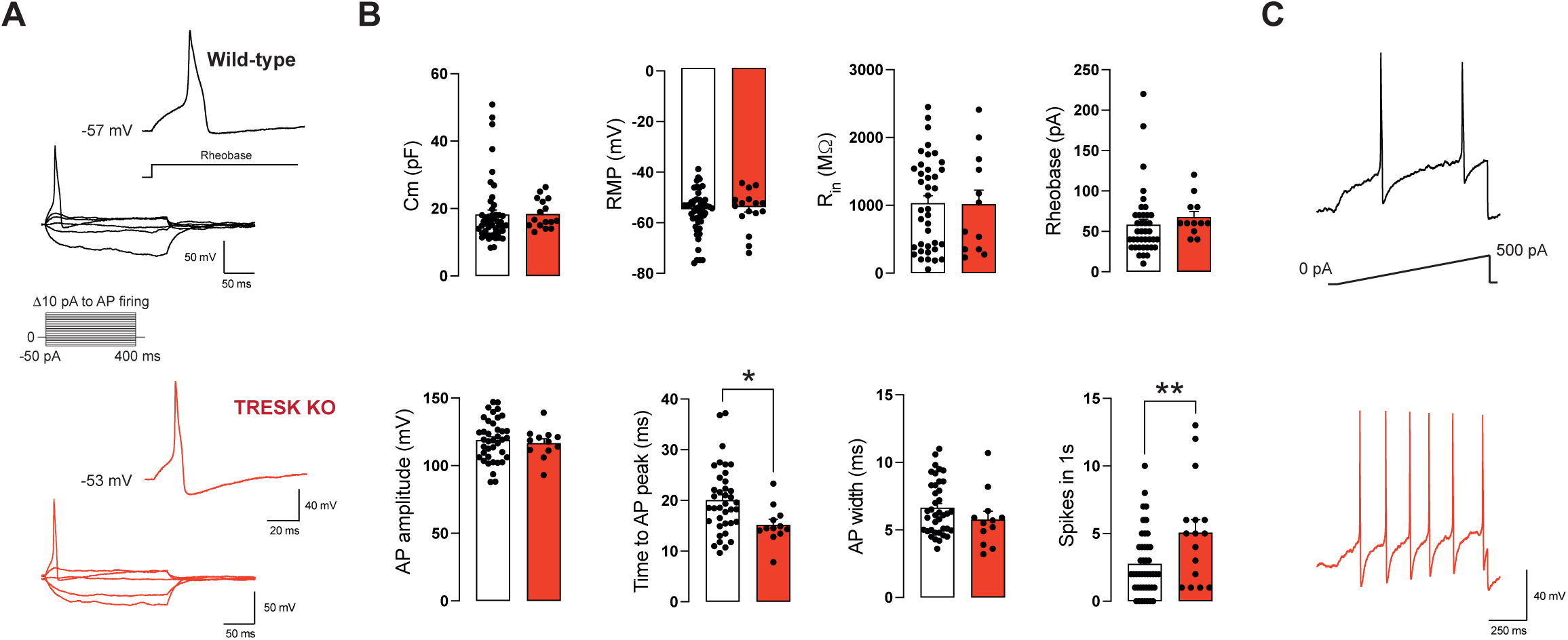
MrgprA3^+^ sensory neurons lacking TRESK exhibit increased excitability. **A.** Representative whole-cell current-clamp recordings from primary cultures of MrgprA3^+^ sensory neurons obtained from in MrgprA3^tdTomato^; TRESK^+/+^ (*left, top*) and MrgprA3^tdTomato^; TRESK^-/-^ (*left, bottom*) mice. Hyperpolarizing or depolarizing 400 ms current pulses in 10 pA increments from −50 pA to action potential firing (Rheobase) were used. For clarity, not all current pulses are displayed. *Inset:* Magnification of the action potential elicited at rheobase. **B.** Quantification of the electrophysiological parameters analyzed in MrgprA3^+^ sensory neurons from wild-type (black bars, n=49) and TRESK^-/-^ (red bars, n=16) animals. Mean membrane capacitance (C_m_) and resting membrane potential (RMP) from neurons recorded is shown. Whole-cell input resistance (R_in_) was calculated from the −50 pA current pulse. Rheobase was measured using 400 ms depolarizing current pulses in 10 pA increments. AP amplitude was measured from the RMP to the AP peak. Time to AP peak was measured from the beginning of the depolarizing pulse to AP peak. AP duration/width was measured at 50% of the AP amplitude. Neuronal excitability was measured as the number of action potential fired in response to a depolarizing ramp from 0 to 500 pA in 1s from RPM. Data is presented as mean±SEM. Statistical differences between groups are shown (*p<0.05, **p<0.01 unpaired two-tailed Student’s t-test). **C.** Examples of neuronal excitability as the number of action potentials fired in response to a depolarizing current ramp (black, wild-type; red, TRESK^-/-^).

### TRESK delays and reduces chronic itching

In addition to systemic and neuropathic conditions, several dermatological conditions contribute significantly to itch, including psoriasis, dry skin (xerosis) and atopic or allergic dermatitis (44,45). As TRESK appears to modulate some forms of acute itch and MrgprA3-expressing neurons have been involved in chronic itch (39,46), we aimed to investigate whether the channel shapes the excitability of sensory afferents during chronic itch. To address this, we initially implemented a well-described mouse model of psoriasiform dermatitis induced by imiquimod (IMQ) treatment, which closely resembles the human disease (37,47). Repetitive application of IMQ in the cheek skin over seven consecutive days developed a psoriasis-like lesion accompanied by chronic spontaneous itch in WT animals. While both male and female mice exhibited evident skin lesion and spontaneous itching, the number of scratching bouts was more pronounced in male mice, with female mice displaying a milder phenotype, consistent with previous reports (48). Consequently, subsequent studies using the IMQ model focused on male mice due to their more robust response, with data from female mice is provided in supplementary Figure 1.

Imiquimod treatment in male WT mice resulted in a progressive increase of spontaneous scratching bouts (Day 2: 14.3±3; Day 3: 34.6±8; Day 4: 54.8±13; p<0.01 vs. Day 0) concurrent with the development of the skin lesion. Persistent itching remained elevated until the conclusion of the protocol (Days 5-7: 41-45 scratching bouts, Fig 4A). Vehicle-treated mice (vaseline) barely exhibited any significant itch response (scratching bouts ranged between 1 to 5; Fig 4A), consistent with observations in basal conditions without any treatment. As expected, the IMQ-treated area displayed a characteristic psoriasis-like skin inflammation, characterized by thickening, scaliness and erythema (37). A different cohort of mice (n=5 mice per group) was used to collect histological samples from cheek skin and mRNA from ipsilateral trigeminal ganglion at day 4 of the treatment (maximum scratching time point). Histological sections from cheek skin showed epidermal hyperplasia with a 3-fold increase in the number of skin epidermal layers compared to vehicle treatment (6.4±0.2 *vs.* 1.9±0.2, respectively; p<0.0001; Fig 4B), as well as hypergranulosis (% epidermal lenght: 7.8±3.4% *vs.* 34.5±5.3%; p<0.05) and a higher number of microabscesses (number/cm: 0.0±0.0 *vs.* 5.1±0.8; p<0.0001). In animals lacking TRESK, a similar skin lesion was observed after IMQ treatment, and no significant differences were observed in histological sections when comparing WT and KO animals treated with IMQ (skin epidermal layers: 6.3±0.2; hypergranulosis: 53.0±6.5%; microabscesses: 3.5±0.6; Fig 4B). Nevertheless, in TRESK^-/-^ mice spontaneous itch was consistently higher compared to wild type animals (ANOVA p<0.0001; IMQ-treated WT *vs.* KO). Spontaneous itch developed earlier in the model and a larger number of scratching bouts were observed (Day 2: 29.6±5.5 scratching bouts; Day 3: 97.0±16.0, p<0.05; Day 4: 141.0±21.8, p<0.05; Fig 4A) compared to IMQ-treated wild-type mice. Later, spontaneous scratching reached similar levels to those found in WT mice, suggesting that the absence of the channel promotes an earlier development of persistent spontaneous itch by facilitating pruriceptor activation and/or enhancing their excitability.

**Figure 4:**
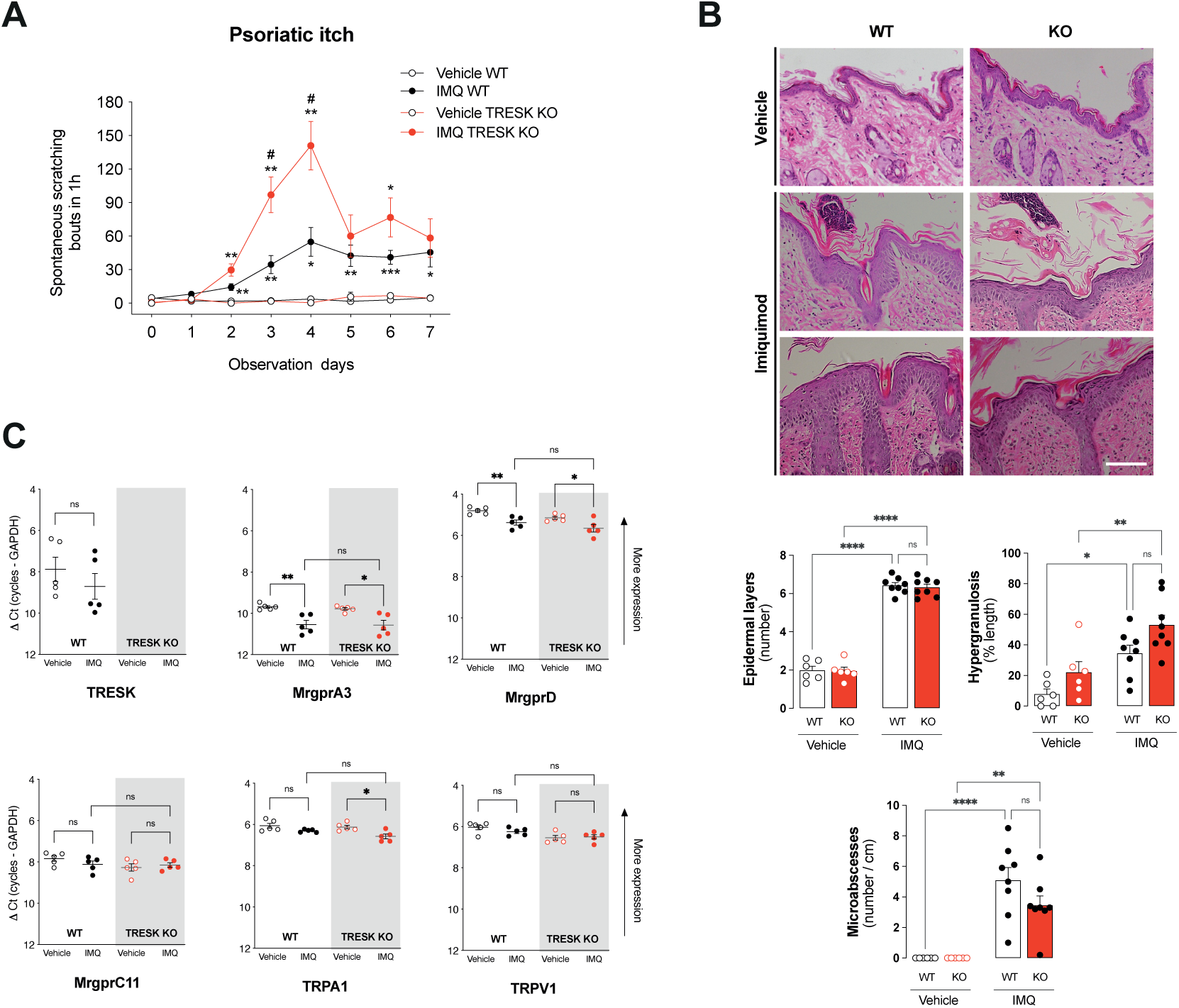
Psoriatic chronic itch is enhanced in the absence of TRESK. **A.** Time-dependent scratching bouts in the IMQ-induced psoriasiform dermatitis model directed to the cheek skin from WT (vehicle, n=8; IMQ, n=15) and TRESK^-/-^ male mice (vehicle, n=4; IMQ, n=8). Significant differences between groups are indicated as: *p<0.05; **p<0.01; ***p<0.001 *vs.* vehicle WT/KO; ^#^p<0.05 IMQ WT vs. IMQ KO; Two-way ANOVA followed by Bonferroni’s multiple comparisons tests. **B.** *Top*. Imiquimod treatment on the cheek skin causes psoriasis-like skin lesions. Representative histological sections stained with hematoxylin and eosin from cheek skin on day 4 of treatment with vehicle (vaseline) or IMQ in WT or TRESK^-/-^ mice are shown. *Bottom*. Quantification of epidermal hyperplasia (as number of epidermal layers), hypergranulosis (as % epidermal length with thickened stratum granulosum) and microabscesses (numbers/cm) in cheek skin samples from WT and TRESK^-/-^ mice. n=6 (vehicle) and 8 (IMQ) per genotype. Scale bar=100 μm. *p<0.05; **p<0.01; ****p<0.0001 *vs.* vehicle WT/KO; One-way ANOVA followed by Holm-Šídák’s multiple comparisons tests. **C.** mRNA expression of TRESK, MrgprA3, MrgprD, MrgprC11, TRPA1 and TRPV1 in trigeminal sensory neurons from WT and TRESK^-/-^ mice at day 4 of IMQ treatment. Data obtained by quantitative PCR shows a significant decrease of MrgprA3 and MrgprD expression in WT and KO, and TRPA1 in KO animals, when comparing with vehicle-treated animals. However, no significant differences were observed between IMQ-treated WT and KO animals. Y-axis shows the ΔCt (number of cycles target - number of cycles GADPH) for the different mRNAs. Notice that the Y-axis has been inverted to visually show that lower ΔCt numbers are indicative of a higher expression. Each dot represents a single animal (n=5 ganglia per group and genotype). Statistical differences between groups are shown (*p<0.05, **p<0.01 unpaired two-tailed Student’s t-test).

Trigeminal ganglion mRNA from the treated side (day 4) was extracted and the expression of different membrane receptors and channels involved in itch detection and transduction was compared in the different groups (Fig 4C). IMQ-treatment did not induce a significant change in TRESK expression. However, both MrgprA3 and MrgprD showed a significant down-regulation on their expression levels, both in WT and KO animals (Fig 4C), suggesting that the skin lesion resulting from IMQ treatment may be inducing a deregulation of these membrane receptors. Similar effects on these receptors have been previously reported in DRGs after IMQ treatment in the back of WT animals (37). No significant changes in expression were observed for MrgprC11, TRPA1 or TRPV1, except a slight but significant decrease in TRPA1 expression in KO animals (Fig 4C).

To further investigate the potential role of TRESK in itch associated with different dermatological conditions, we conducted a comparative analysis between WT and KO animals in two additional skin diseases models, allergic contact dermatitis (ACD) and dry skin (Fig 5). In the ACD model, animals were initially sensitized through topical application of SABDE on the back skin and subsequently challenged with the compound in the mouse cheek a few days later, following stablished protocols (39). After the first challenge, WT animals showed an increased number of scratching bouts directed to the treated area compared to vehicle or to values obtained previously to allergen application (Vehicle: 1.9±0.6 scratching bouts; Pre-challenge: 1.3±0.5; 1st challenge: 18.4±3.0; p<0.001 *vs.* pre-challenge; Fig 5A). Similar itching scores were observed during the second challenge with the allergen one day later (18.1±3.3; p<0.01 *vs.* pre-challenge values), indicating that allergen sensitization resulted in a persistent effect upon each subsequent challenge with the allergic compound. In this model, KO animals displayed a more pronounced effect on the scratching score on the first day of the challenge (Vehicle: 2.6±0.7 scratching bouts; Pre-challenge: 1.6±0.7; 1st challenge: 30.5±4.5; p<0.001 *vs.* pre-challenge; p<0.05 *vs.* WT mice; Fig 5A), suggesting a enhanced activation of peripheral sensory afferents. On the second challenge, both WT and KO did not differ in the number of scratching bouts elicited (2nd challenge WT: 18.1±3.3; KO: 21.7±3.9), despite both groups presented significant itching compared to vehicle application.

**Figure 5:**
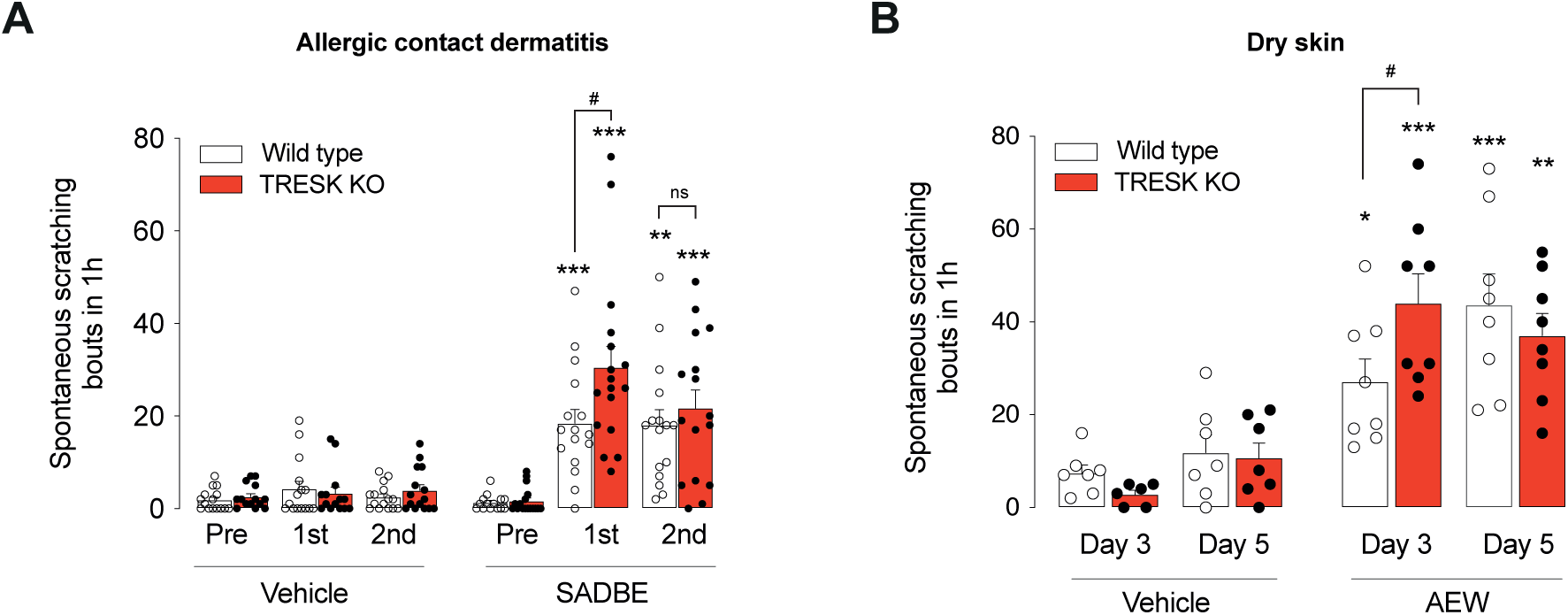
TRESK deletion also enhances chronic itch in other models of skin disease. **A.** Spontaneous scratching bouts in the allergic contact dermatitis model. Animals were initially sensitized with SADBE topical application in the back skin and then challenged with the compound in the mouse cheek a few days later. The number of scratchings directed to the cheek during 1h are plotted before (Pre), 1 day after (1st) and 2 days after (2nd) repetitive SADBE cheek application. WT (vehicle, n=15; SADBE, n=16) and TRESK^-/-^ mice (vehicle, n=15; SADBE, n=17). Significant differences between groups are shown as: **p<0.01; ***p<0.001 vs. vehicle WT/KO Two-way ANOVA followed by Bonferroni’s multiple comparisons tests.; ^#^p<0.05 SADBE WT vs. KO; Two-way ANOVA followed by Bonferroni’s multiple comparisons tests. **B.** Spontaneous scratching bouts in the dry skin model. After repetitive application of acetone, ether and water (AEW) to the cheek skin, the number of spontaneous scratching bouts during 1h on day 3 and 5 are plotted. WT (vehicle, n=7; AEW, n=8) and TRESK^-/-^ mice (vehicle, n=7; AEW, n=8). Significant differences between groups are shown as: *p<0.05; **p<0.01; ***p<0.001 vs. vehicle WT/KO ^#^p<0.05 AEW WT vs. KO Two-way ANOVA followed by Bonferroni’s multiple comparisons tests.

A similar pattern was observed in the dry skin model, where repetitive application of acetone, ether, and water (AEW) in the cheek led to the development of a skin lesion that induced robust itch behaviors on day 3 and beyond (38). As anticipated and compared to the vehicle (7.4±1.7 scratching bouts), WT mice presented a substantial spontaneous pruritus on day 3 (27.1±4.9; p<0.05 *vs.* vehicle; Fig 5B), which further intensified at day 5 of treatment (43.6±6.8; p<0.001 *vs.* vehicle). Similarly, to the ACD model, scratching induced by dry skin manifested earlier in the absence of TRESK channel, with maximum scratching values observed as earlier as day 3 (44.0±6.4; p<0.001 *vs.* vehicle and p<0.05 *vs.* WT; Fig 5B) and remaining consistent on day 5 (37.0±4.8; p<0.01 *vs.* vehicle; ns *vs.* WT). The similar pattern of early itch development across the three skin models suggests that the presence of the channel impedes or delays the development of persistent pruriceptor activation and in its absence, sensory neurons display an enhanced susceptibility to activation by pruritic agonists.

### Pharmacological activation of TRESK reduces itch

In contrast to histaminergic itch, which has a plethora of antihistaminergic drugs available, the ability to modulate certain forms of non-histaminergic itch has long been pursued due to the lack of effective treatments or specific pharmacological compounds. Given that the absence of TRESK correlates with an enhancement of acute and chronic itch, both for therapeutic purposes and to further demonstrate the role of this channel, we evaluated whether its activation diminish scratching behavior, serving as a functional readout of itch sensitivity. Cloxyquin (Clx), an old anti-bacterial and anti-fungal drug used in tuberculosis treatment, has been identified as specific TRESK opener among the other K_2P_ channels (49,50), with an effect independent of the Ca^2+^/calcineurin-mediated potentiation of the channel current. Initially, we examined whether cloxyquin could prevent chloroquine-induced acute itch in the cheek model. Intraperitoneal injection of cloxyquin (50 mg/Kg) or its vehicle (olive oil), administered two hours before the pruritic test did not induce any significant scratching in wild-type mice (Fig 6A). In line with previous observations (Fig 2A), chloroquine injection resulted in abundant scratching bouts directed to the cheek (61.4±3.9; p<0.0001 *vs.* vehicle). Conversely, in animals previously injected with cloxyquin, a significant reduction (57%) in the number of scratching bouts was obtained (35.1±2.4; p<0.0001 CQ+ Clx *vs.* CQ+veh; Fig 6A), indicating that TRESK activation greatly reduced the number of observed scratchings. To confirm that the effect of cloxyquin was due to TRESK channel activation and to rule out other potential side effects, we conducted the same experiment in mice lacking the channel. As previously demonstrated, in TRESK^-/-^ mice, chloroquine produced a significantly larger effect compared to WT animals (79.6±8.2 scratching bouts *vs.* 61.4±3.9; p<0.05; Fig 6A), and this effect was not altered after cloxyquin administration (74.4±5.8; p>0.99 *vs.* CQ+veh). The significant difference compared to WT mice (p<0.0001) indicates that the channel must be present to obtain the therapeutic effect.

**Figure 6:**
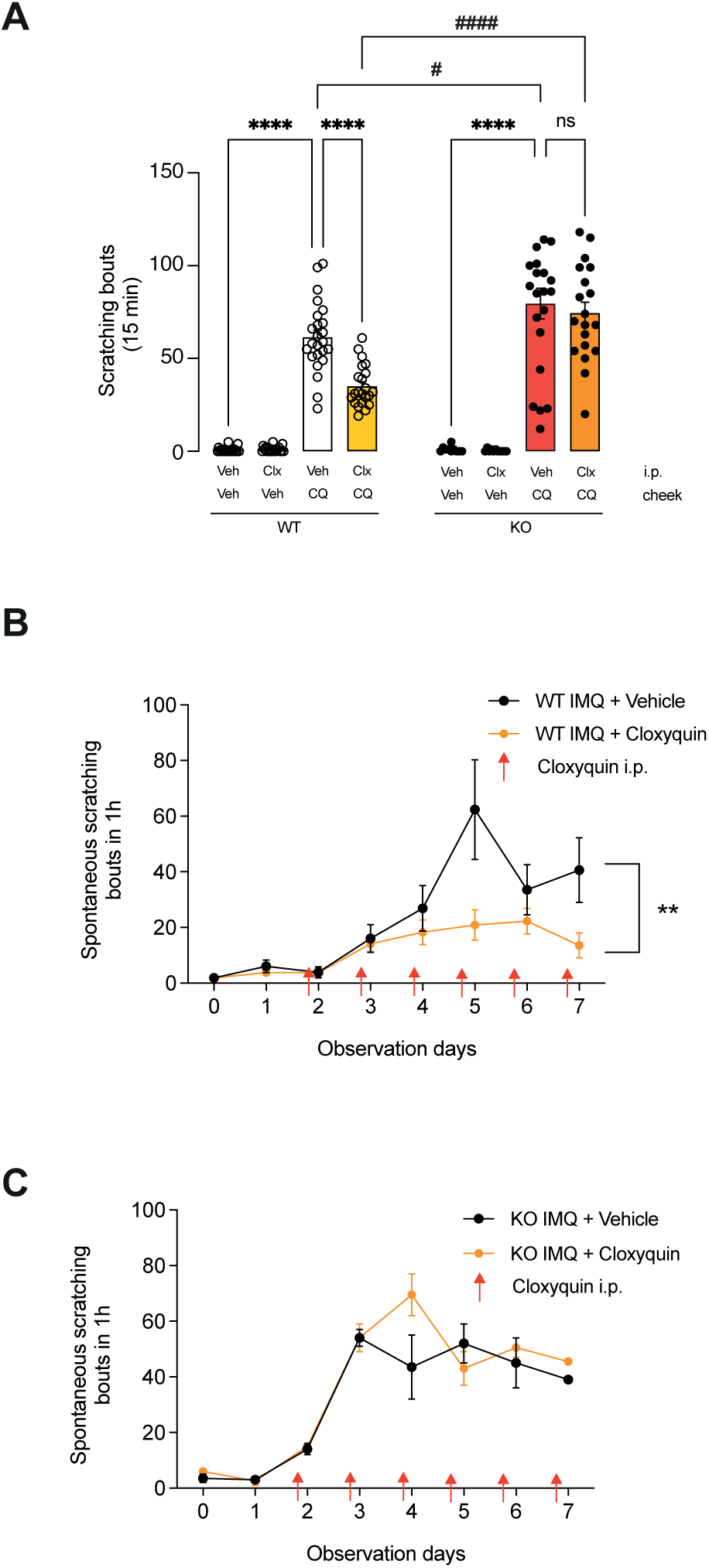
Pharmacological activation of TRESK reduces acute and chronic itch. **A.** Acute itch produced by intradermal injection of chloroquine (CQ; 50 μg) into the cheek was measured during 15 min, 2h after i.p. injection of cloxyquin (Clx; 50 mg/Kg) or its vehicle (olive oil). Neither vehicle (olive oil) nor cloxyquin produced significant scratching bouts with vehicle (PBS) cheek injection. CQ-induced scratchings were greatly reduced by pretreatment with Clx. The reducing effect of Clx was not observed in TRESK^-/-^ animals, which corroborates the specific TRESK- mediated effect of the drug in WT animals. WT (vehicle/vehicle, n=21; vehicle/Clx, n=21; vehicle/CQ, n=24; Clx/CQ, n=22) and TRESK^-/-^ mice (vehicle/vehicle, n=10; vehicle/Clx, n=11; vehicle/CQ, n=21; Clx/CQ, n=19). Significant differences between groups of the same genotype are shown as: ****p<0.0001. Differences between WT and KO groups are shown as ^#^p<0.05 and ^####^ p<0.0001. Two-way ANOVA followed by Bonferroni’s multiple comparisons tests. **B.** Time-dependent scratching bouts in the IMQ-induced psoriasiform dermatitis model in the cheek skin from WT male mice. Mice were injected with cloxyquin (Clx; 50 mg/Kg; n=9) or vehicle (olive oil; n=6) 2h before observation as indicated by red arrows. A significant difference between treatments is shown as **p<0.01 Two-way ANOVA. **C.** Time-dependent scratching bouts in the imiquimod-induced psoriasiform dermatitis model in the cheek skin from KO male mice. Mice were injected with cloxyquin (Clx; 50 mg/Kg; n=4) or vehicle (olive oil; n=4) 2h before observation as indicated by red arrows. No statistically significant differences were obtained.

We next investigated whether the beneficial effect of TRESK activation extended to a model of chronic itch. Using the IMQ-induced of psoriatic itch model, we administered cloxyquin or vehicle (i.p.) once daily from days 2 to 7, 2 hours before assessing scratching behavior. Vehicle injection yielded no significant effect, while, as anticipated, IMQ treatment in male mice led to a progressive increase in the number of scratchings events (Fig 6B). In contrast, cloxyquin treatment notably diminished spontaneous scratching over several consecutive days (p<0.01; Fig 6B), once again underscoring that the modulation of TRESK reduces the development of spontaneous itch in this model. As a control, TRESK^-/-^ mice developed IMQ-induced psoriatic itch, but no effect of the channel activator was observed (Fig 6C).

## Discussion

Different populations of sensory neurons have been identified as pruriceptors, characterized by the specific expression of receptors for pruritogenic compounds. Besides the effects of histamine through the activation of its receptors (histaminergic pathway), numerous other receptors activate non-histaminergic pathways that convey multiple pruritic signals through the spinal cord or the brain stem towards the brain. The initial step in this pathway is the activation of peripheral afferents. As primary sensory neurons undergo activation, the likelihood that the depolarization triggered by a stimulus reaches action potential threshold depends on the coordinated activity of several ion channels, encompassing different types of Na^+^, K^+^ and Cl^-^ channels (51). Channels active in the subthreshold range of membrane voltages, situated between resting membrane potential and action potential threshold, play a crucial role. Among these channels, K_2P_ channels function as a break preventing neuronal activation and/or excessive neuronal excitability (52). In this context, we and others have described that down-regulation of TRESK, after injury, inflammation or bone metastasis contribute to enhanced responsiveness of sensory neurons and resultant hyperexcitability, leading to persistent pain (22,53–56). We have previously shown that knocking out TRESK, which is prominently expressed in sensory neurons, enhances the excitability of these cells, and increases the sensitivity to mechanical and cold pain (24). Data extracted through comprehensive transcriptomic studies have shown significant TRESK expression in non-peptidergic NP1 and NP2 subpopulations of mouse sensory neurons and in pruritogen receptor-enriched neurons in humans (4,13,21,29–32,57,58). Our present data confirm a significant co-expression of TRESK with MrgprD and MrgprA3 receptors, two well-established markers for NP1 and NP2 neuronal subtypes in mice, respectively. This specific expression pattern suggests that TRESK is well-placed to specifically regulate the excitability of these neuronal subtypes. Functionally, we observed that acute skin injection of chloroquine, the MrgprA3 receptor agonist, elicited abundant scratching behavior directed to the site of injection, effect that was more pronounced in TRESK^-/-^ animals compared to WT ones. This observation suggests that the channel plays a regulatory role in the excitability of NP2 neurons. To pharmacologically dissect this effect, different pruritogens were used to activate selected receptors in specific pruriceptor subpopulations. Histamine receptor Hrh1 is predominantly expressed in MrgprA3^+^ (NP2) and Nppb^+^ (NP3) neurons (59). Injection of histamine did not render any significant difference in scratching number between WT and KO animals, suggesting that a combined effect in all these populations masks any enhanced response due to TRESK ablation. Indeed, after ablation of MrgprA3^+^ neurons with diphtheria toxin (5), mice still present a scratching response to histamine, indicating that sensitivity to histamine is not exclusively mediated by MrgprA3^+^ neurons. In fact, the TRESK has a minimal expression in NP3 neurons and likely plays a minor role in their excitability (4,21). To corroborate that TRESK does not regulate the excitability of NP3 neurons, we used Leukotriene C4 (LTC4), an agonist of Cysltr2 receptor specifically expressed in Nppb^+^ neurons (59). As expected, we did not observe any difference in the scratching response in animals expressing or lacking TRESK, confirming that the channel does not have a major role preventing their activation. MrgprA3^+^ neurons have been further subdivided into NP2.1 and NP2.2 (4). To distinguish between both subpopulations, we used BAM8-22, whose receptor MrgprC11 (or MrgprX1) is exclusively expressed in NP2.2 neurons (4,21). Despite NP2.2 neurons expressing MrgprA3 and TRESK at lower levels compared to NP2.1 neurons, the channel seems to play a minor role in these cells, as BAM8-22 induced similar scratching responses in WT and TRESK^-/-^ animals. Therefore, our data indicates that the enhanced response to chloroquine is mainly mediated by activation of NP2.1 neurons. Similar percentages of neurons were activated by chloroquine in calcium imaging experiments in WT and KO animals, ruling out that the enhanced effect in KO animals is due to a larger population of MrgprA3^+^ neurons activated by this compound. Likewise, other pruritogens activated similar percentages of neurons, indicating that TRESK removal in KO animals does not appear to produce any remodeling of pruriceptor populations.

Different mechanisms contribute to the transduction of pruritic signals into electrical signals in sensory neurons. It has been reported that TRPA1 acts downstream of MrgprA3 activation, and it is required for chloroquine-evoked itch (12). A Gβγ signaling mechanism couples MrgprA3 to TRPA1, leading to extracellular Ca^2+^ entry, while Ca^2+^ release from intracellular stores is not required, as occurs with MrgprC11-evoked itch (12). However, conflicting evidence arises from skin nerve preparation studies, where the activation of cutaneous C fibers by chloroquine remains unaffected in the presence of a TRPA1 blocker or in *Trpa1*-deficient mice but requires the participation of phospholipase Cβ3 (60). Another report has shown that the Ca^2+^-dependent chloride channel ANO1 also plays a role in transmitting Mrgpr-dependent itch in the periphery, as itch mediated by chloroquine, SLIGRL or bilirubin (Mrgpr-dependent pruritogens) is reduced in Ano1-deficient mice (16). These pruritogens appear to activate ANO1 downstream of their membrane receptor, leading to neuronal depolarization. In vivo experiments also demonstrate that chloroquine activation of sensory neurons can be significantly blocked by a ANO1 blocker (16). The precise mechanism for MrgprA3 receptor signaling to activate sensory neurons remains to be fully elucidated, with potential involvement of TRPA1, ANO1 or other mechanisms such as TRPC3 (61). Interestingly, both TRPA1 and ANO1 expression appear to be low in MrgprA3^+^ sensory neurons, while ANO3 presents a higher expression (21,46,61). Our findings reveal that TRESK contributes to MrgprA3-dependent itch, likely by counteracting the depolarization induced by activation of TRPA1 and/or ANO1. Beyond membrane depolarization, calcium entry into the sensory neuron will further enhance TRESK current by calcineurin-dependent dephosphorylation of the serine cluster in the intracellular loop of the channel (62). Therefore, TRESK, in conjunction with other hyperpolarizing mechanisms, modulates the neuronal firing and, consequently, itch sensitivity. Our experiments validate the initial prediction that the absence of TRESK facilitates neuronal firing and enhances scratching behaviors following chloroquine skin injection. Some reports highlight the multimodal capacity of some sensory neurons, like MrgprA3^+^ ones, which express different receptors in the same cell type (e.g. MrgprA3, MrgprC11, TRPV1), but activate different signal transduction mechanisms to trigger pain or itch (17). It is plausible that TRESK activity is specifically linked to some of these mechanisms, but no others, as TRESK-deficient mice exhibit enhanced itch in response to chloroquine stimulation but not to BAM8-22. Both compounds activate membrane receptors in the same cell type but likely initiate different signal transduction pathways (12,60).

The mechanisms governing the excitability of MrgprA3^+^ cells are not well-understood, and the contribution of different ion channels expressed has not been precisely defined, as has been in other neuronal subtypes. In these subtypes, the distinctive expression of Na^+^, K^+^, Ca^2+^ and Cl^-^ ion channels is a key determinant of subtype-specific intrinsic physiological properties (19). Electrophysiological characteristics of MrgprA3^+^ neurons recorded in the present study align with those described previously (63). In comparison to other sensory neurons (MrgprA3^-^), MrgprA3^+^ population exhibit a lower action potential frequency in response to depolarizing current injection. We have observed a similar pattern, and repetitive firing is mostly absent to suprathreshold stimuli injection. However, MrgprA3^+^ neurons from TRESK-deficient mice present an increased firing upon current ramp injection, akin to what has been previously reported in nociceptive neurons (24). This observation suggests once again that TRESK plays a role in modulating the excitability of these neurons.

MrgprA3^+^ neurons play a significant role in the pathophysiology of chronic itch. Deletion of the Mrgpr-cluster or specific ablation of MrgprA3^+^ neurons has been shown to decrease itch in dry skin, contact dermatitis, and allergic dermatitis mouse models (5,64). Additionally, induction of chronic dermatitis through skin treatment with SADBE increases the excitability of MrgprA3^+^ neurons (39). In the AEW model of dry skin, has been observed an increase in the number of MrgprA3^+^ peripheral fibers, DRG neurons and MrgprA3 mRNA (65). In the present study, we have shown that in both the allergic dermatitis and the dry skin model, the absence of TRESK enhances the number of spontaneous scratching behaviors, suggesting that increased activation of MrgprA3^+^ neurons facilitate the development of the disease.

MrgprA3^+^ neurons seem to also mediate the effects of endogenous pruritogens such as peptide GPR15L, which is increased in the skin of patients with psoriasis and atopic dermatitis (66). GPR15L is released from inflamed keratinocytes and activates different Mrgpr receptors in mast cells and sensory neurons, including MrgprA3 in mice and MrgprX1 and X3 in humans (66). Psoriasis, a chronic skin inflammatory disease characterized by hyperplasia, hyperkeratosis and severe chronic itch (67). Repeated topical application of imiquimod, a Toll-like receptor 7 agonist, serves as a commonly used mouse model for the disease, inducing psoriatic skin lesions and itch that closely resembles the human condition, particularly in the B6 mouse strain (37,68). Our data indicate that mice lacking TRESK exhibit an earlier onset and an significant increase in the number of scratches, suggesting that the absence of the channel promotes neuronal activation and development of the itching associated psoriatic phenotype. As observed previously (48), imiquimod skin treatment led to a substantial increase in spontaneous scratching bouts in WT males, but only a minor increase in females, despite similar skin lesion and transcriptional IMQ signatures in both sexes (48,68). In our current study, we noted that TRESK-deficient mice displayed even higher spontaneous itch behavior throughout the entire mouse model Although females exhibited a small number of spontaneous scratching bouts, all KO animals present higher spontaneous itch responses, albeit the differences were much smaller compared to the IMQ-induced phenotype in males (Fig.4A).

Despite not observing significant differences in the skin lesion between WT and KO male mice at day 4 (when the maximum of scratching bouts was observed), it is possible that enhanced itch in KO animals facilitates an earlier development of the skin lesion. In this context, hypergranulosis seems slightly increased in the KO mice (p=0.107) and it cannot be ruled out that at late time points (days 5 to 7) some differences in skin pathogenic parameters exist. Whether an earlier development of the skin lesion in KO mice occurs due to an enhanced neuronal activation or due to the more important mechanical effect remains to be elucidated. Possible differences may exist at earlier time points (days 2 or 3), but it appears that the IMQ-induced psoriasiform phenotype fully develops similarly in both genotypes. It is also conceivable that a continuously elevated number of scratches worsen the skin lesions in the long run, and enhancing TRESK activation might ameliorate the disease. Furthermore, both WT and KO mice used in this study have a C57Bl/6N background, which exhibits a milder phenotype in the IMQ-induced psoriasiform dermatitis model compared to the same model in C57Bl/6J mice, likely due to lower expression of IL-22 in the former (69). Any difference in the skin lesion between WT and KO mice could potentially be observed in C57Bl/6J mice, which presents a more aggravated dermatitis.

At the genetic level, the IMQ-induced psoriasiform phenotype significantly regulates a multitude of genes, both in the skin and in sensory neurons innervating the affected area, mirroring the processes observed in human psoriasis (37,68,70). We did not observe significant differences in the mRNA expression of *trpa1, trpv1* or *mrgprc11* in trigeminal ganglia, when comparing IMQ-treated animals form both genotypes. These findings suggest that the expression of these channels and receptors is not significantly altered in the model. However, it also raises the possibility that potential gene expression changes in neurons innervating the affected area might be masked by the overall expression of all neurons in the trigeminal ganglia. To conclusively confirm potential expression changes, specific gene expression evaluation at a single-cell level in neurons innervating the affected area must be conducted. Although no differences between genotypes were observed in IMQ-treated animals, as previously reported (37), we noted a mRNA downregulation of MrgprA3 and MrgprD in both genotypes, and TRPA1 only in KO animals, when comparing vehicle and IMQ-treated animals. This observation could be explained by a downregulation due to persistent activation. Regarding TRESK, we did not observe any significant difference in WT animals when comparing vehicle and IMQ-treated animals. Assuming that there are no gene expression changes and without validation at a single cell level, this observation opens up the possibility of using TRESK as a therapeutic target in chronic itch. Moreover, part of the pruritogenic effect of IMQ in sensory neurons has been attributed to inhibitory effects on some Kv and K_2P_ channels (71). We also observed an inhibitory effect of IMQ in TRESK (data not shown), as reported for TREK-1 and TRAAK. This inhibitory effect can contribute to chronic effects of IMQ, in addition to the immune and inflammatory mechanisms involved. The fact that IMQ-induced skin histological phenotype is similar in WT and TRESK^-/-^ animals indicates that TRESK is not critically involved in the development of IMQ-induced keratinocyte proliferation, differentiation, and inflammatory response. The absence of TRESK results a similar phenotypic pattern in the other models of chronic itch, with enhanced pruritus during the initial days of the model (day 3 in dry skin, 1st challenge in allergic contact dermatitis). This implies that TRESK serves to limit sensory neurons activation when present, although these differences fade as the disease progresses.

To explore therapeutic potential, we investigated whether the TRESK activator cloxyquin could improve the scratching phenotype by hampering sensory neuron activation. Our data shows that pretreatment with cloxyquin to enhance channel activation significantly diminishes the number of scratching episodes in the chloroquine injection model of acute itch in WT, but not in the KO mice (Fig 6A), implying a specific effect of cloxyquin on TRESK channels. Moreover, although we only tested a single cloxyquin concentration applied 2h before the pruritic challenge, the data demonstrates that TRESK activation can be employed to alleviate pruritus and further underscore the role of this channel modulating pruriceptor excitability. Similarly, in the IMQ model of chronic itch, pretreatment of animals (2h before behavioral assessment) with cloxyquin from day 2 (once the spontaneous itch behaviors become apparent), also attenuated the pruritic phenotype in WT mice, while this effect was not observed in KO mice. Although refining the therapeutic dose and administration pattern may enhance pruritus inhibition, these experiments highlight the potential development of TRESK activators as antipruritic agents for non-histaminergic itch, either alone or in combination with other drugs targeting different mechanisms (e.g. Mrgpr inhibitors).

In summary, we describe here that TRESK background potassium channel regulates the excitability of specific subset of pruriceptors, and its absence enhance both acute and chronic itch. Enhancing the channel function with specific activators constitutes a novel anti-pruritic therapeutic approach that can be combined with other compounds for the treatment of non-histaminergic itch, for which appropriate treatments are lacking.

## Figure Legends

**Supplementary Figure 1: Psoriatic chronic itch model in female mice** Time-dependent scratching bouts in the IMQ-induced psoriasiform dermatitis model directed to the cheek skin from WT (n=7) and TRESK-/- female mice (n=5). Despite an increase in the number of scratching bouts over days of treatment, no significant differences were found between groups. Notice the reduced number of scratching bouts compared to males (Fig 4A), which indicates that female mice display a milder phenotype, as reported by others (see ref. 48).

## Authors’ contributions

Authors AAB, GC and XG performed electrophysiological recordings in neurons and calcium imaging. JLlA, AAB, IP and NC performed behavioral experiments. AAB, IP and GC carried out primary cell cultures. AAB, JLlA and APC performed in situ hybridization and qPCR experiments. CS, JMdA and AAB performed the histological experiments and analysis. JLlA, AAB, CS, NC, GC and XG participated in the design of the study and performed the statistical analysis. GC and XG conceived the study, oversaw the research and prepared the manuscript with help from all others. All authors read and approved the final manuscript.

