## Supplementary figures and images for "TRESK background potassium channel in MrgprA3^+^ pruriceptors regulates acute and chronic itch"

### Supplemental Fig 1

Psoriatic itch - Females

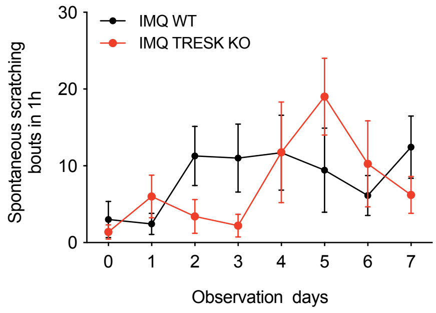


#### Supp Fig 1
